# DeepGAMI: Deep biologically guided auxiliary learning for multimodal integration and imputation to improve phenotype prediction

**DOI:** 10.1101/2022.08.16.504101

**Authors:** Pramod Bharadwaj Chandrashekar, Jiebiao Wang, Gabriel E. Hoffman, Chenfeng He, Ting Jin, Sayali Alatkar, Saniya Khullar, Jaroslav Bendl, John F. Fullard, Panagiotis Roussos, Daifeng Wang

## Abstract

Genotype-phenotype association is found in many biological systems, such as brain-related diseases and behavioral traits. Despite the recent improvement in the prediction of phenotypes from genotypes, they can be further improved and explainability of these predictions remains challenging, primarily due to complex underlying molecular and cellular mechanisms. Emerging multimodal data enables studying such mechanisms at different scales from genotype to phenotypes involving intermediate phenotypes like gene expression. However, due to the black-box nature of many machine learning techniques, it is challenging to integrate these multi-modalities and interpret the biological insights in prediction, especially when some modality is missing. Biological knowledge has recently been incorporated into machine learning modeling to help understand the reasoning behind the choices made by these models.

To this end, we developed DeepGAMI, an interpretable deep learning model to improve genotype-phenotype prediction from multimodal data. DeepGAMI uses prior biological knowledge to define the neural network architecture. Notably, it embeds an auxiliary-learning layer for cross-modal imputation while training the model from multimodal data. Using this pre-trained layer, we can impute latent features of additional modalities and thus enable predicting phenotypes from a single modality only. Finally, the model uses integrated gradient to prioritize multimodal features and links for phenotypes. We applied DeepGAMI to multiple emerging multimodal datasets: (1) population-level genotype and bulk-tissue gene expression data for predicting schizophrenia, (2) population-level genotype and gene expression data for predicting clinical phenotypes in Alzheimer’s Disease, (3) gene expression and electrophysiological data of single neuronal cells in the mouse visual cortex, and (4) cell-type gene expression and genotype data for predicting schizophrenia. We found that DeepGAMI outperforms existing state-of-the-art methods and provides a profound understanding of gene regulatory mechanisms from genotype to phenotype, especially at cellular resolution. DeepGAMI is an open-source tool and is available at https://github.com/daifengwanglab/DeepGAMI.

## Introduction

Genotype-phenotype association has been found in many biological systems such as brain-related diseases and behavioral traits. However, predicting phenotypes from genotypes remains challenging, primarily due to complex underlying molecular and cellular mechanisms. Genome-wide association studies (GWAS) investigate genotype-phenotype associations using genotyping microarray and whole-genome sequencing techniques. A typical GWAS determines the association of genetic variants with heritable diseases^1,2^. GWAS discoveries have led to the discovery of numerous disease-associated variants. More than a hundred GWAS studies have been conducted on neurodegenerative and psychiatric diseases like Alzheimer’s disease (AD)^3–6^ and schizophrenia (SCZ)^7,8^, among other brain diseases. Despite the ground-breaking findings from these GWAS studies, they have some limitations. Firstly, association studies do not imply causation and require further downstream analysis and validations. Secondly, GWAS studies are independent studies that try to find the relationship between variants and disease individually and fail to tackle the combined effect. Finally, the SNPs having small effect sizes go undetected as they do not meet the threshold criteria of the existing studies^9^. There have been several computational attempts outside the GWAS studies to discover genotype-phenotype association. Most of these attempts involve regression^10–12^, with Polygenic Risk Scores^13^ being the most popular and widely used method that looks at the linear combined effect of several variants on the phenotype. Modern machine learning techniques like Convolution Neural network (CNN) have been applied to predict the functionality of these phenotypes. For example, Zhang et al.^14^ grouped several SNPs based on LD structure and used it as an input to the CNN model to predict AD progression. These approaches consider genotype to phenotype prediction and fail to address the complex nature of this association that mainly involves a lot of intermediate phenotypes like gene expression and epigenome regulation.

Several studies have shown that these variants influence disease risks by altering cell-type regulatory elements that in-turn affect the underlying gene expressions that affect the disease phenotype^15,16^. This resulted in studying the effects of genotypes on gene expression. Expression quantitative trait loci (eQTL) studies focus on associating genetic variants to gene expression instead of disease phenotypes^17–22^. They have proved to be a critical step in investigating regulatory pathways and have identified numerous eQTLs that modulate the expression of genes associated with various human diseases. However, these studies fail to determine the causality of the variants. Transcriptome-wide association studies (TWAS) aim at identifying gene-trait interaction by combining GWAS and gene expression. The effect of genetic variation on gene expression is first studied, and then these expression profiles are associated with the traits using statistical methods^23–30^. A significant drawback in TWAS is that co-expressed gene patterns often lead to many false hits and may prioritize non-causal genes^31^. PrediXcan^32^ is a popular computational approach that predicts gene expression levels from eQTLs and then maps trait-associated loci based on the predicted gene expression levels. TWAS and PrediXcan perform association between the traits and the gene expression. Association studies suffer from the causality issue meaning that it requires further additional downstream techniques to identify causal genes.

It is also necessary to analyze genes and other regulatory elements that impact disease phenotypes. Several computational approaches have attempted to associate genes with disease risks using gene expression profiles directly. For example, Taesic Lee and Hyunju Lee^33^ collected gene expression profiles from three publicly available Alzheimer’s (AD) datasets to predict the onset of AD disease using a variational autoencoder (VAE). DeepWAS^25^ first predicted regulatory functionality (like histone modification and chromatin interaction) of the genetic variants using DeepSEA^35^ and then applied regression to predict the phenotype. A different approach is to look at gene regulatory networks (GRNs). GRNs are inferred by gene expression data representing a group of genes interacting with each other to control several biological mechanisms. They have proven helpful in mapping molecular interactions^36,37^ and biomarkers for brain diseases^38–40^. A key point to note here is that these GRNs are cell-type specific. A major drawback of these approaches is that they consider each omics individually for modeling the relation between the omics and the disease phenotype (like genotype to genotype or gene expression to disease).

As the biological processes involve a complex interaction with multi-omics, emerging multimodal data enables studying such mechanisms at different scales. Several studies have attempted integrating multi-omics data for brain disease predictions like SCZ^41–43^ and AD^44,45^. Most of these existing approaches for integrating multimodal data do not consider known biological findings (like GRNs, and eQTLs) for guiding these predictive models. Some studies have successfully used biological knowledge to help guide feature selection or integrate several omics for disease prediction. For example, Wang et al.^46^ uses a deep Boltzmann machine (DBM) architecture for genotype to phenotype prediction, where gene regulatory networks guide the internal connections for predicting SCZ, BPD, and ASD diseases. Varmole^47^ integrated gene expression and genotype (SNPs) using a dense neural network architecture where GRN and eQTLs guide the relationship between the input and the first hidden layer. Most of these studies consider disease outcomes as the phenotypes that try to predict disease versus control. However, more complex phenotypes, like neuropathological phenotypes (Braak score, global cognitive function, plaque count) for AD and cellular phenotypes like cell layers in the mouse visual cortex, need to be studied. Considering these different phenotypes for associating with the genotypes can help us understand the disease’s finer cellular, molecular, and pathological mechanisms. Another critical factor is the availability of multimodal data. Many studies extract only one type of modality or partially extract multimodal data. Due to several factors (experimental bottlenecks, sample availability, etc.), muti-omics data will often be partially available for a given individual^48^. For example, given a particular cohort, one might find genotype information but lacks (or contains a small number of) gene expression, ATAC-seq, or imaging data^49,50^. In these cases, it will be difficult to include these partially available datasets into a predictive model due to the lack of enough samples and missing data. This calls for cross-modal data imputation that can be further used for prediction. No existing approaches perform cross-modality estimation alongside predicting disease phenotypes to the best of our knowledge. Auxiliary learning has been gaining attention to improve the generalization of the primary task^51–54^. An auxiliary task can be considered as a subtask that can be trained along with the primary task where the features are shared between the tasks. Auxiliary tasks are usually defined in estimating entities relevant to solving the main task and were initially referred to as hints^55^. One of the widely used ways of implementing auxiliary learning is adding supplementary cost to the primary cost function of the neural network model^56^. Although no existing approaches use auxiliary learning for data imputation, the closest one is SCENA^57^ which tries to estimate the gene-gene correlation matrix using ensemble learning and auxiliary information for single-cell RNA-seq (scRNA-seq) data.

To this end, we propose DeepGAMI: <u>Deep</u> biologically <u>g</u>uided <u>a</u>uxiliary learning for <u>multimodal i</u>ntegration and imputation to improve phenotype prediction. DeepGAMI is a novel deep learning approach that integrates genotype and gene expression data for predicting disease phenotypes using auxiliary learning of cross-modality estimation. Our contributions are three-fold. Firstly, we present a deep learning framework that integrates genotype and gene expression data guided by prior biological knowledge of QTLs and GRNs. Secondly, with the help of auxiliary learning, our framework can take in only one data modality, perform cross-modal estimation by learning linear relationships between different modalities, and use the estimated values for disease prediction. Thirdly, we decipher the black-box nature of the neural network architecture to identify and prioritize genes and SNPs contributing to disease onset. We applied DeepGAMI to multiple emerging multimodal datasets, including population-level genotype and bulk and cell-type gene expression data for SCZ cohort^46^, genotype and gene expression data for AD cohort^50^, and recent single-cell multimodal data comprising transcriptomics and electrophysiology for neuronal cells in the mouse visual cortex^58^. We found that DeepGAMI outperforms existing approaches in predicting phenotypes among all three datasets and also prioritizes genes, SNPs and electrophysiological features having biological significance and relevance. DeepGAMI is an open-source tool and is available at https://github.com/daifengwanglab/DeepGAMI.

## Results

### DeepGAMI overview

DeepGAMI is a multi-view interpretable deep learning model that integrates multimodal data for predicting phenotypes. The model architecture of DeepGAMI is shown in **Fig 1**. Importantly, it uses an auxiliary learning mechanism to learn the latent space of one modality from another, thereby enabling us to predict phenotypes from the available modalities by performing cross-modality imputation. The model also uses biological prior to aid in prediction and further guide us in interpretability in the form of prioritizing genes, SNPs, and other biological features. We denote the primary task as 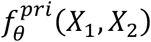 that takes available multimodal inputs *X*_1_ and *X*_2_ with parameters *θ* to predict phenotypes, e.g., *X*_1_ and *X*_2_ can be SNP genotypes and gene expression levels of individuals. The auxiliary task is denoted by 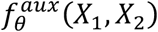 which aims at learning *C*_*X*_2__ (latent space of input *X*_2_) from *C*_*X*_1__ (latent space of input *X*_1_). Using the predictive model learnt by available multimodal inputs, DeepGAMI can predict the phenotypes of the samples with single modality only. Specifically, it first imputes other modal latent spaces using the auxiliary learning function and then feeds both imputed and input latent spaces into the primary task for phenotype prediction.

**Figure 1:**
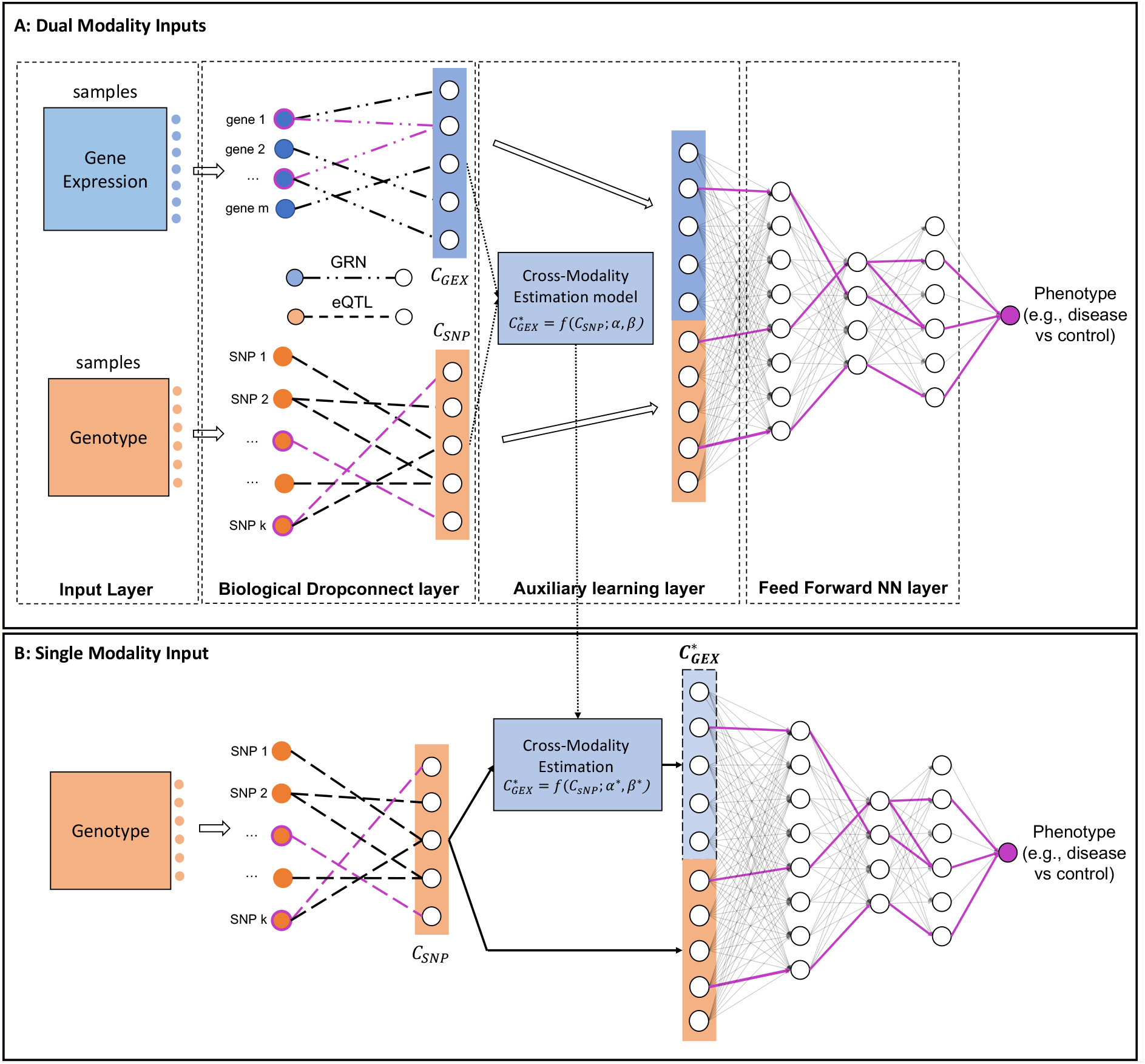
Architecture of DeepGAMI. (**A**) DeepGAMI first uses available multimodal features for training the predictive model, e.g., SNP genotypes (orange) and gene expression (blue) of individuals from the major applications in this study. In particular, it learns the latent space of each modality (e.g., consisting of latent features at the first transparent hidden layer). This learning step is also regularized by prior knowledge enabling biological interpretability after prediction, i.e., the input and latent features are connected by biological networks (biological dropConnect). For instance, the input transcription factor genes can be connected to target genes as their latent features (e.g., *C_GEX_*) by a gene regulatory network (GRN). The input SNPs can be connected to associated genes as their latent features (e.g., *C_SNP_*) by Expression quantitative trait loci (eQTLs). Notably, an auxiliary learning layer is used to infer the latent space of one modality to another, i.e., cross-modality imputation. For instance, DeepGAMI learns a transfer function *f*(.) to estimate *C_GEX_* from *C_SNP_*. Finally, the latent features are concatenated and fed to the feed-forward neural networks for phenotype predictions, e.g., classifying disease vs. control individuals. (**B**) Using the learned predictive model from multimodal input along, DeepGAMI can predict phenotypes from a single modality, e.g., SNP genotypes of new individuals. Specifically, it first imputes other modal latent spaces using the optimal transfer function *f*\*(.) and then feeds both imputed and input latent features into downstream neural network predictions, e.g., *C*^*^_*GEX*_ from *C_SNP_*.

### Classification of schizophrenia individuals from genotype and bulk and cell-type gene expression data

We first tested DeepGAMI on population-level genotype and bulk-tissue gene expression data of individuals (dorsolateral prefrontal cortex or DLPFC brain region). In particular, we used the population data from the PsychENCODE consortium^46^ for predicting schizophrenia (SCZ) versus healthy individuals. After filtering and preprocessing (Methods and Materials), we ended up with 2080 SNPs, 126 TFs, and 84 intermediate layer genes for classifying 343 control and 275 SCZ individuals.

**Fig 2A and Fig 2B** shows the five-fold cross-validation balanced accuracies and AUC scores respectively of DeepGAMI (Dual modality where both genotype and gene expression are given to the model as input and Single modality where only genotype is provided as an input) in comparison to all the baseline models (Random Forest, Naïve Bayes, MLP), Varmole, and DeepGAMI with no biological priors. DeepGAMI dual (*BACC* = 0.845 ± 0.022) and DeepGAMI single (*BACC* = 0.831 ± 0.024) outperform all the baseline and state-of-the-art models. DeepGAMI with no biological prior (*BACC* = 0.827 ± 0.0 for dual modalities and *BACC* = 0.789 ± 0.019 for single modality) is the closest. DeepGAMI dual (AUC = 0.923) and DeepGAMI single (AUC = 0.878) also performs better in comparison to PRS scores (AUC = 0.61)^59^. Incorporating biological priors into the model has the advantage of better interpreting SNPs, genes, and regulatory networks. **Fig 2C** shows a few examples of prioritized SNPs and genes along with prioritized functional links by the Integrated Gradient method (Methods and materials). Our model can prioritize SCZ-related genes like GAD1^60,61^, MCC^62^, and TOMIL2^63^ shown as pink circles in **Fig 2C**. A complete list of prioritized SNPs and genes with the importance score is available in **Supplementary Data S1 and S2**. We used the prioritized functional links to extract the genes present in these links (top-ranked 10% of the link importance scores) and performed enrichment analysis on them. We found several known functions and pathways related to SCZ, like response to oxidative stress^64–66^, membrane trafficking^67^, and, GPCR signaling^68,69^, etc.

**Figure 2:**
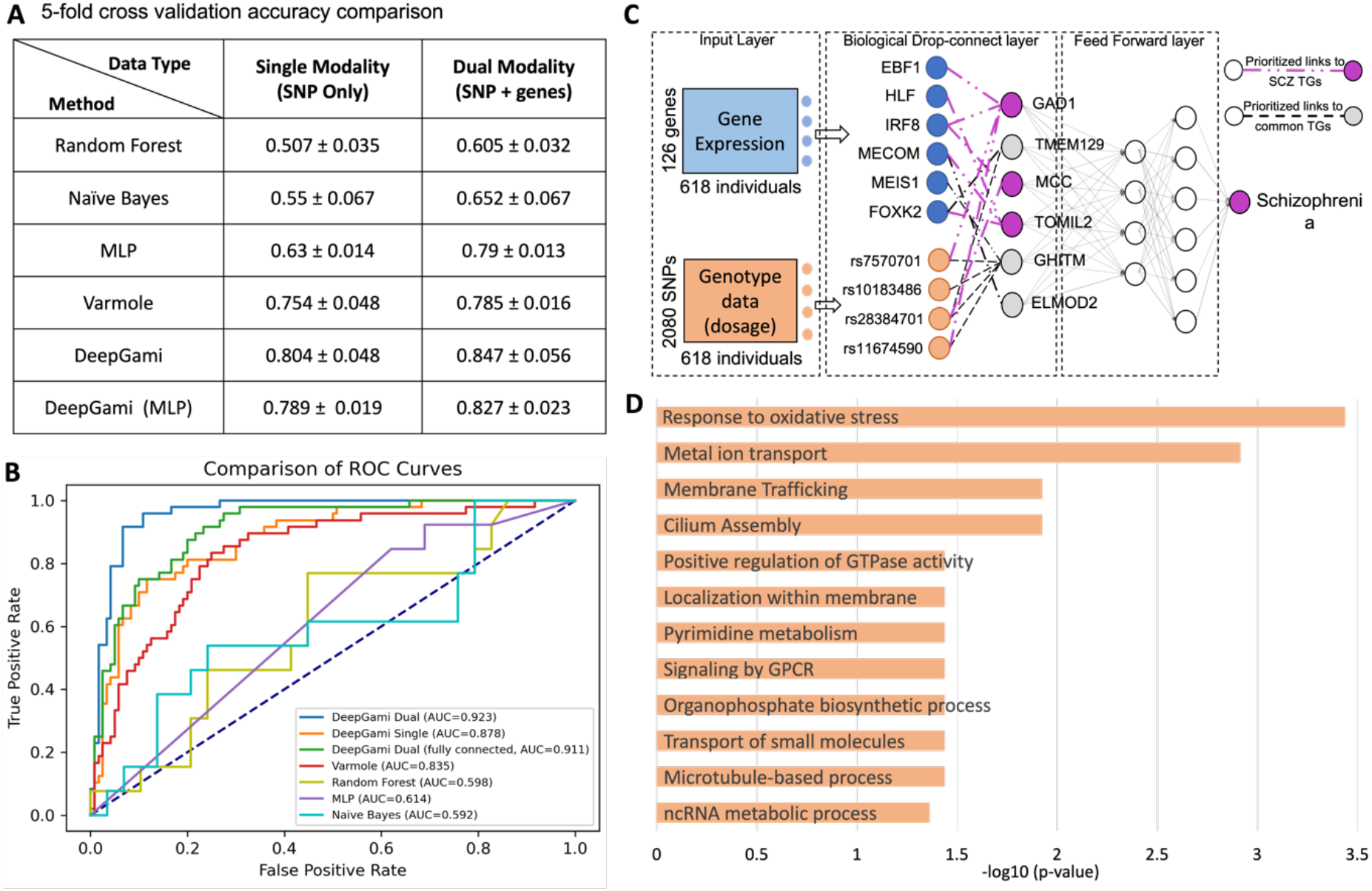
Schizophrenia classification and functional genomic prioritization using genotype and bulk-tissue gene expression data. The population data was from the PsychENCODE project (Methods and Materials). (**A**) Balanced Accuracies from 5-fold cross-validation and **(B)** Receiver operating characteristic curves of DeepGAMI dual-modality model (dark blue), DeepGAMI single modality model (orange), DeepGAMI without any biological priors (green, i.e., fully connected) and the state-of-the-art algorithms: Naïve Bayes (light blue), Random Forest (yellow), Multilayer perceptron (MLP, purple), Varmole (Red) for classifying schizophrenia vs. control individuals. **(C)** Select examples of prioritized transcription factors, SNPs, target genes (latent features, and functional links (GRNs, eQTLs) for schizophrenia. Purple: known schizophrenia genes. **(D)** Function and pathway enrichments of prioritized schizophrenia SNPs.

### Clinical phenotype prediction and gene regulatory network prioritization in Alzheimer’s disease

To demonstrate the application of DeepGAMI for predicting complex clinical phenotypes, we ran DeepGAMI on an Alzheimer’s disease (AD) cohort ROSMAP^50^. ROSMAP contains multi-omics data of the DLPFC brain region for aging and AD among humans from Rush University. We use the bulk-tissue gene expression and genotype data from this cohort for our analysis. We extracted the eQTL information from the GTEx consortium^70^ for the human brain frontal cortex (BA9), which contains 146,763 eQTL SNPs, and used the GRN from the psychENCODE consortium^46^. Clinical phenotypes included in our analysis are cognitive diagnosis (COGDX), CERAD, and BRAAK scores. **Supplementary Table 2** summarizes the number of features and class labels used for this analysis.

We split the data into training and held-out test sets using an 80:20 ratio. We used a five-fold CV on the training set for tuning the hyperparameters and obtaining the best performance. DeepGAMI outperforms state-of-the-art classifiers (BACC = 0.806 for BRAAK, 0.689 for COGDX, and 0681 for CERAD, **Fig 3A, Supplementary Table 3**). For BRAAK phenotype, DeepGAMI (*BACC* = 0.806 ± 0.03 for dual, *BACC* = 0.79 ± 0.02 for single) outperforms random guess (BACC = 0.50), Random Forest classifier(*BACC* = 0.538 ± 0.01), Naïve Bayes classifier (*BACC* = 0.555 ± 0.03), and MLP (*BACC* = 0.742 ± 0.02). Looking at a more complex phenotype with multi-class, DeepGAMI with two modality inputs improved the classification accuracy of COGDX by 1.72 folds (*BACC* = 0.688 ± 0.07) compared to the highest accuracy of the baseline model (Naïve Bayes, *BACC* = 0.399 ± 0.05). Also, DeepGAMI with single modality input improved the multi-class classification accuracy by 1.71 folds ((*BACC* = 0.686 ± 0.07)) compared to the Naïve Bayes classifier (*BACC* = 0.399 ± 0.05).

**Figure 3:**
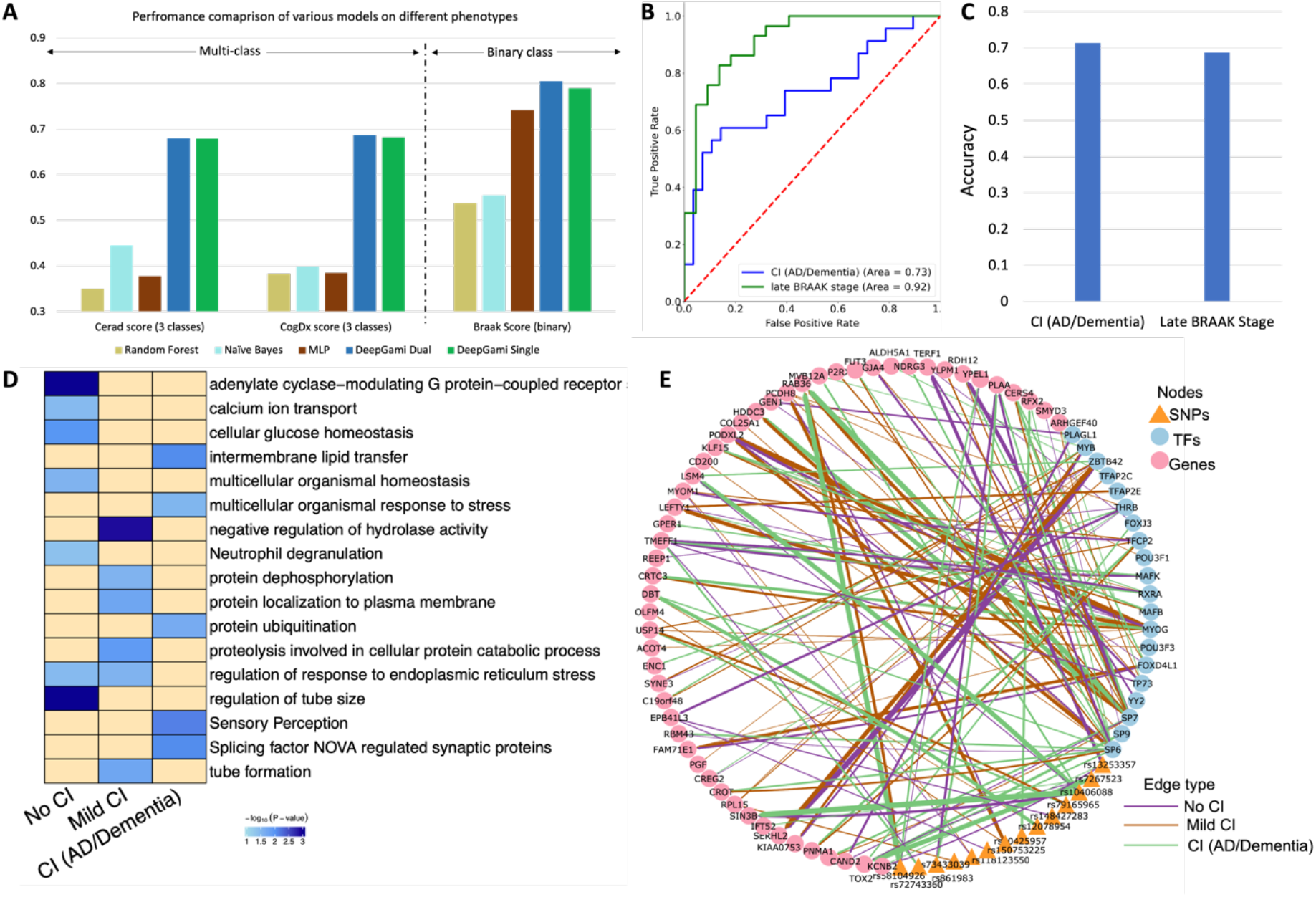
Multi-class clinical phenotype prediction and regulatory network prioritization in Alzheimer’s disease. **(A)** 5-fold cross-validation performance of DeepGAMI (Dual modality: dark blue, Single modality: green) on three different phenotypes: neuritic plaque measure (CERAD score, multi-class), cognitive impairment (COGDX score, multi-class) and neurofibrillary tangle pathology (BRAAK stage, binary) in comparison with Naïve Bayes (light blue), Random Forest (yellow), MLP (brown). **(B)** ROC curves of held-out test samples for cognitive COGDX phenotype (blue) and late BRAAK stage (green). **(C)** Classification accuracies of the independent dataset for COGDX phenotype and BRAAK stage. **(D)** Enrichment analysis of prioritized genes for no cognitive impairment, mild cognitive impairment, and cognitive impairment (AD/Dementia) classes of COGDX phenotype. **(E)** Select an example of a prioritized regulatory network for the cognitive impairment phenotype. The edge thickness between any two nodes corresponds to the prioritized link importance score of the associated nodes. The edge color represents the three classes.

As our goal was to predict phenotypes with missing modalities (single modality as input), we tested DeepGAMI to classify the held-out test data containing late-stage BRAAK individuals and CI(AD/Dementia) COGDX phenotypes using only genotype as input. DeepGAMI was able to classify these samples accurately (**Fig 3B**) with late-stage BRAAK AUC of 0.92 and CI(AD/Dementia) COGDX AUC of 0.73. Furthermore, ROSMAP contains additional individuals with only genotype data. We were able to classify late-stage BRAAK individuals with an accuracy of 0.687 accuracy and CI(AD/Dementia) COGDX individuals with an accuracy of 0.713 (**Fig 3C**).

We then performed enrichment analysis on the top-ranked genes that were regulated by the SNPs and TFs for each group (No CI, mild CI, CI(AD/Dementia)) of the COGDX phenotype (**Fig 3D, Supplementary Data S3**) and generated a network containing the prioritized SNPs, TFs, with the prioritized links to the genes (**Fig 3E, Supplementary Data S4**). We found that these prioritized genes are enriched with many known cognitive impairment functions and pathways. For example, controls were enriched for adenylate cyclase-modulating G protein-coupled receptors that are known to have a role in pathological prognosis of AD^71,72^. Mild CI was associated with protein dephosphorylation^73^, repones to endoplasmic reticulum stress^74–76^ and proteolysis in cellular protein catabolic process^77^. We observed that CI is associated with sensory perception and splicing factor NOVA regulated synaptic proteins. Sensory perception impairment is known to affect cognition^78–80^. NOVA regulates genes critical for neuronal function^81^ and known to affect patients with inhibitory motor control dysfunctions^82^.

### Cortical layer classification for single-cell neuronal cells in mouse visual cortex

To test if DeepGAMI can perform well using additional data modalities of single cells, we ran DeepGAMI on an emerging Patch-seq dataset^58^ containing single-cell multimodal data for the visual cortex brain region in neuronal cells of mouse species. This dataset includes transcriptomics and electrophysiological (ephys) data. We used the cell layers (L1, L2/3, L4, L5, and L6) that reveal the location of the cells in the visual cortex as the cellular phenotype. We followed the data extraction and preprocessing as done in DeepManReg^83^ and ended up with 41 ephys features and 1000 genes for 3,654 cells.

We deviated from our five-fold cross-validation splits and instead, we followed the approaches used by existing approaches and randomly split the cells into training/testing sets with a ratio of 80:20 and obtained 100 different sets. For each training set, we applied oversampling to have a balanced number of cells for each layer in the training label and ended up with 941 cells in each layer. Biological Dropconnect was not used as there was no biological prior data available. We instead used full connectivity, where each gene and ephys feature had an association with the intermediate latent space layer. After various parameter tuning, the latent space dimension was set to 500. We used the Kolmogorov-Smirnov test (k.s test) to compare the prediction accuracy across different methods. DeepGAMI has higher prediction accuracy for classifying cell layers than other methods (k.s test statistic=1, *p* – value < 2.2*e*^−16^ for dual-modality input, k.s test statistic=1, *p* – *value* < 1.9e^−16^ for single modality gene expression input, **Fig 4B** and **Supplementary Table 4**). Furthermore, average accuracy of DeepGAMI dual-modality mode (65.71% with a 95% confidence interval (CI) of [62.49%, 68.92%]) and DeepGAMI single modality mode (64.63% [61.29, 67.97]) is higher than the random guess baseline of 20% (five labels), LMA^84^ (43.0% [32.2%, 49.6%]), CCA^85^ - 46.2% [40.1%, 51.3%], MATCHER^86^ - 46.5% [40.9%, 52.8%]), and DeepManReg^83^ - 51.4% mean accuracy, [47.9%, 54.8%])).

**Figure 4:**
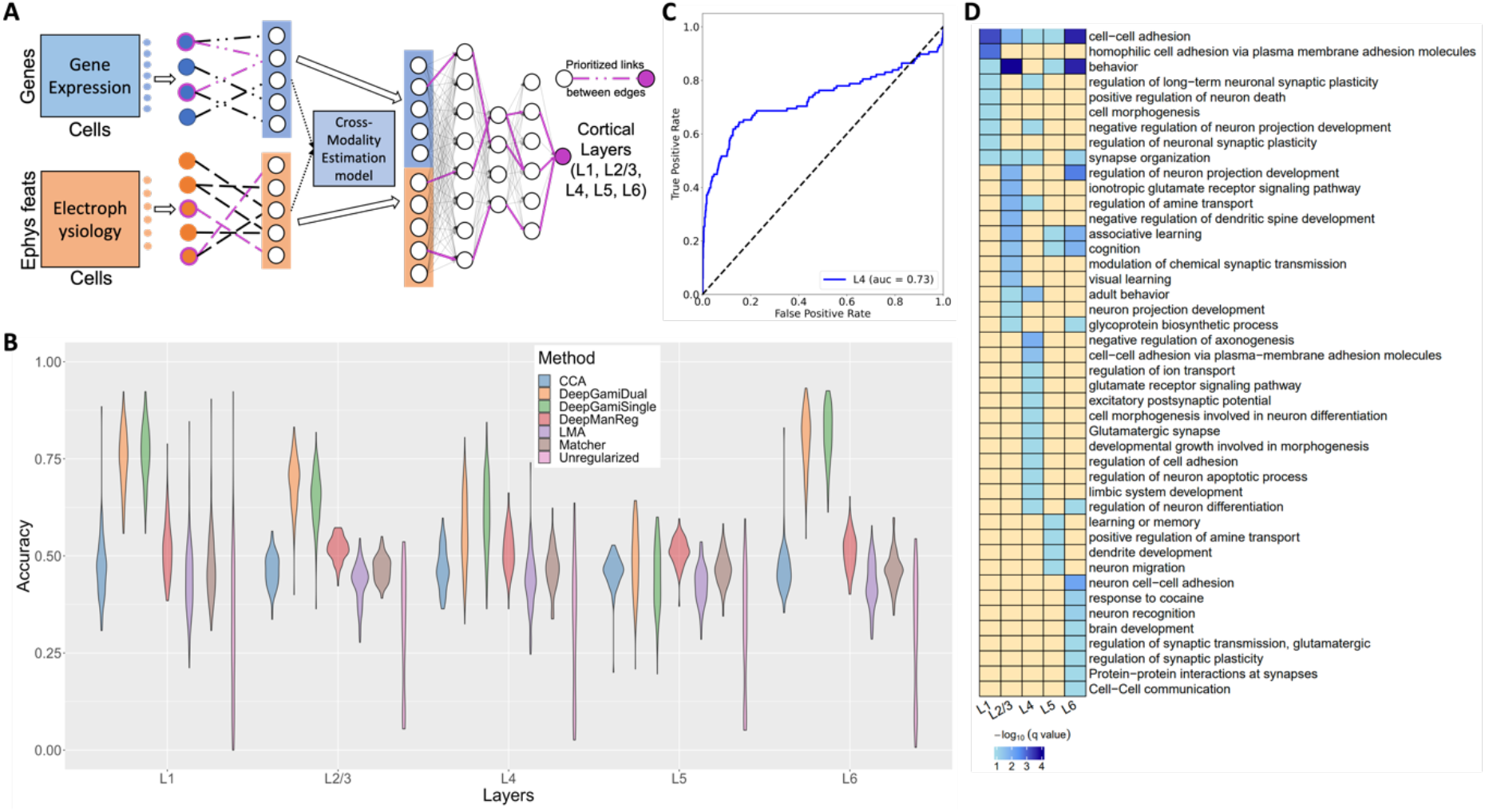
Classifying cellular phenotype in single neuronal cells of mouse visual cortex. **(A)** DeepGAMI model for cell layer classification. **(B)** Balanced accuracies for classifying cell layers in the mouse visual cortex by DeepGAMI dual-modality (orange), DeepgGAMI single-modality (green) versus DeepManReg^83^ (dark pink), neural network classification without any regularization (light pink), LMA^84^ (violet), CCA^85^ (blue), and MATCHEr^86^ (brown). **(C)** ROC curve for classifying independent L4 visual cortex data. **(D)** Gene enrichment analysis showing the enriched terms for layer-specific prioritized genes.

We then extracted 112 (L4) neuronal cells patch-seq data^87^ containing gene expression data but only a small set of electrophysiological data for independent testing. We applied DeepGAMI for classifying these 112 cells by giving only gene expression (single modality mode) and allowing the model to estimate the latent space of ephys features. For the negative samples required for performance estimation, we used the predictions on the motor cortex data^58^. The motor cortex dataset contains 1286 genes and 29 electrophysiological features. The cell layers comprise of four layers: L1, L2/3, L5, and L6. After predicting these cells, we used the predicted values for the L4 layer as the negative samples, combined them with the predictions for the L4 layer for the 112 samples, and computed the AUC score. DeepGAMI classified cells into L4 layer with an AUC score of 0.73 (**Fig 4C)**.

Following the prediction, we extracted the top 10% of the prioritized genes and importance scores of the 41 ephys features for each cell layer (**Supplementary Data 5**). The prioritized genes were given to metascape^88^ for enrichment analysis. **Fig 4D** shows the gene set enriched terms for each layer. Enriched terms like cell-cell adhesion and neuron projection development appear in all layers^89^. Layer 4 is enriched with excitatory neurons and their activities^90,91^. Many groups were enriched for behavior (especially L2 and L6), and synapse organization. L1 and L2/3 groups were enriched for negative regulation of neuron projection development and long-term neuronal synaptic plasticity regulation. The upper layers of the cortex (groups L2/3 and L4) were enriched for amine transport regulation and adult behavior; the primary input from the thalamus goes to Layer 4, whose input then goes to Layers 2 and 3. **Supplementary Figure 1** compares the importance scores of the 41 ephys features across six cell layers.

### Classification of schizophrenia individuals using genotype and cell-type gene expression data

Following the success of DeepGAMI on bulk-tissue and single-cell data, we tested if DeepGAMI can prioritize cell-type genes and SNPs for four major brain cell types: excitatory neurons, inhibitory neurons, oligodendrocytes, and other glia (microglia and astrocytes). In particular, we used genotype and cell-type specific gene expression imputed using bMIND^92^ from a reference panel of 4 cell populations: GABAergic (i.e. inhibitory) neurons, glutamatergic (i.e. excitatory) neurons, oligodendrocytes, and a remaining group composed mainly of microglia and astrocytes. **Supplementary Table 5** summarizes the number of features used as input for the model for these four cell types. DeepGAMI can classify SCZ individuals with a balanced accuracy of 0.795 ± 0.035 for dual-modality mode and 0.784 ± 0.024 for single modality mode in comparison to 0.563 ± 0.049 for Random Forest, 0.633 ±0.05 for naïve Bayes, 0.743 ± 0.058 for MLP, and 0.765 ±0.026 for varmole in mixture of microglia and astrocytes. Similarly, DeepGAMI displays better five-fold classification performance compared to other models for the other three cell types, as seen in **Fig 5A** (**Supplementary Table 6)**.

**Figure 5:**
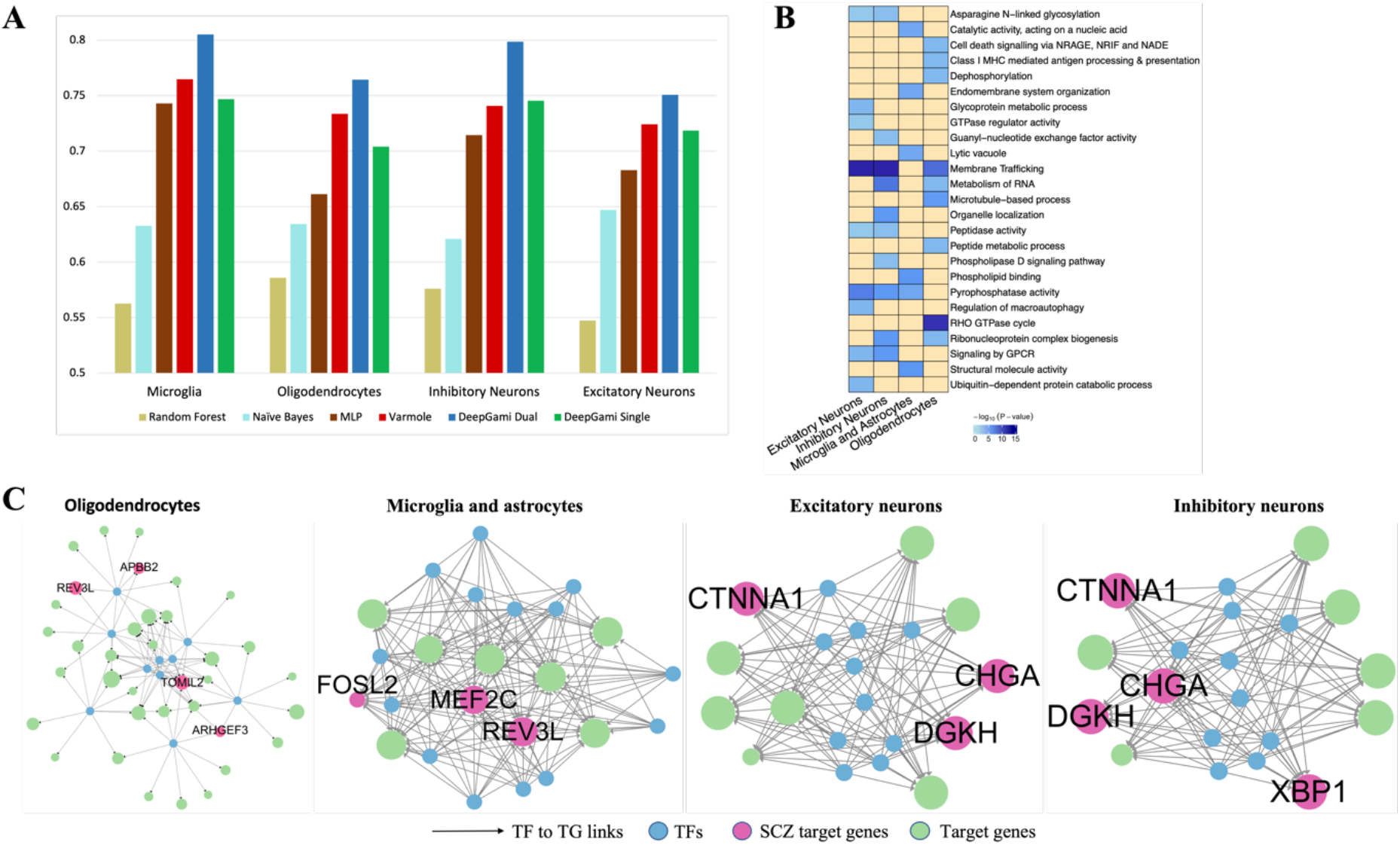
Classification of schizophrenia individuals and prioritization of genes, SNPs, and regulatory network using genotype and gene expressions of four cell types (microglia and astrocyes, oligodendrocytes, inhibitory neurons, and excitatory neurons). **(A)** Balanced accuracies from 5-fold cross-validation of DeepGAMI dual-modality model (dark blue), DeepGAMI single modality model (green) in comparison with Naïve Bayes (light blue), Random Forest (yellow), MLP (brown), and varmole (red). **(B)** Pathway enrichment of prioritized schizophrenia SNPs for four cell types. The blue shade gives the - log(p-values) of the term. **(C)** Prioritized cell-type gene regulatory networks with pink circles representing schizophrenia genes. The size of the target gene is defined by the number of prioritized links between the SNPs and the associated gene.

We then performed enrichment analysis for the prioritized SNPs for each cell type and found various known cell-type pathways and functions associated with SCZ (**Fig 5B**) like organelle localization in inhibitory neurons^93^, structural molecule activity in microglia and astrocytes^94^, RHO GTPase cycle in oligodendrocytes^95^, and regulation of macroautophagy in excitatory neurons^96^. We also found some common SCZ associated functional pathways among multiple cell types like Asparagine N-linked glycosylation^97,98^, membrane trafficking^67^, and signaling by GPCR^69^.

**Fig 5C** represents networks of the prioritized regulatory network for the four cell types. The pink circles represent genes associated with SCZ. The edge weight is proportional to the number of SNPs associated with that particular gene. Our model can prioritize cell-type genes like APBB2 and TOMIL2 for oligodendrocytes, FOSL2 and MEF22C for microglia and astrocytes, and XBP1 for inhibitory neurons, as well as some common genes like REV3L (oligodendrocytes and microglia and astrocytes) and CTNNA1 (excitatory and inhibitory neurons). A complete list of importance scores for every gene and SNPs for each cell type is available in **Supplementary Data S6 and S7**.

## Discussion

In short, DeepGAMI is a novel interpretable deep learning framework for improving genotype-phenotype prediction from multimodal data. It introduces an auxiliary learning layer into the framework to learn the latent features of different modalities from each other. This layer thus enables cross-modal imputation so that DeepGAMI can still predict phenotypes when some modalities are unavailable. The model also takes given biological information as priors for aiding in prioritizing multimodal features (e.g., SNPs, genes) and feature networks (e.g., gene regulatory networks) related to the phenotypes. We show that DeepGAMI can improve phenotype prediction and prioritize phenotypic features and networks in multiple multimodal datasets, including population-level genotype and gene expression data at both tissue- and cell-type levels for brain disorders (e.g., schizophrenia, Alzheimer’s disease), and gene expression and electrophysiological data of single-cell neurons in mouse cortex.

As brain phenotypes typically involve complex cellular and molecular mechanisms, genotype and gene expression are a few of the many factors associated with mechanisms. Integrating additional multi-modalities gives us a more profound understanding of such mechanisms from genotype to phenotype. For example, several studies have tried integrating copy number variations with DNA methylation^99^, gene expression^100^, and clinical data^101^. Trevino et al.^102^ integrated RNA-seq and ATAC-seq over a period of time, studying various genetic activities and disease susceptibility in various neuropsychiatric disorders. Thus, we expect to extend DeepGAMI to more emerging modalities, like epigenetics and imaging, especially in brain research. Moreover, the DeepGAMI framework is currently limited in finding biological insights related to the phenotypes as the model architecture is guided by biological priors about input modalities, e.g., eQTLs and regulatory networks linking SNPs and genes. Having comprehensive data about phenotype-specific biological knowledge may help DeepGAMI perform better in discovering new biological features and novel mechanisms.

The cross-modal imputation in DeepGAMI is carried out by linear regression, assuming that the latent features of different modalities have a linear relationship. Even though our applications here have shown the imputations work very well, the DeepGAMI framework can use nonlinear functions for cross-modality imputation, aiming to have nonlinear auxiliary learning. We mainly focus on integrating two modalities in this work, but we are intrigued to extend DeepGAMI in the near future to include multiple modalities and predict continuous phenotypes. For instance, DeepGAMI opens avenues for integrating several modalities like imaging, epigenomics, behaviors, and clinical records alongside cross-modal imputation to predict complex phenotypes.

## Methods and Materials

### Model design

The DeepGAMI model consists of four main layers.

#### Input layer

The input layer consists of two data modalities, e.g., gene expression and SNP genotypes. The gene expression matrix comprises of *m* genes and *k* samples and is represented by *X_GEX_* ∈ *R^k*m^*. The genotype matrix consists of dosage information of various SNPs (expected allelic counts) *X_SNP_* ∈ *R^k*n^*(*n* SNPs by *k* samples).

#### Biological DropConnect layer

DropConnect is a regularization mechanism that sets random activation units to zero in each layer. It differs from dropout, which sets the random output units to zero while the former sets the connection weights to zero^103^. For our purpose, instead of randomly setting activations to zero, we guide the activations using prior biological knowledge, as shown in **Equations 1 and 2**.

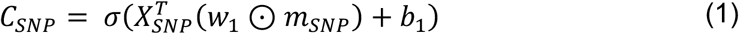

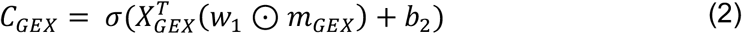

where *w*_1_ *and w*_2_ are weight vectors, *b*_1_ *and b*_2_ are the bias, and ⊙ is the Hadamard product (element-wise multiplication). The input to this layer is the *X* matrix, the output nodes comprise genes, and *m* encodes biological drop connections (**Equations 3 and 4**)

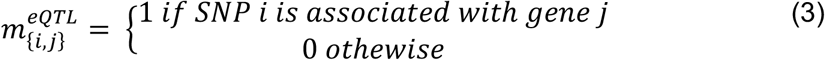

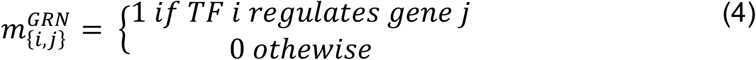

This enables us to understand several mechanisms affecting gene regulation, further guiding us towards better phenotype prediction. We call the output of this layer the latent space of the input matrix.

#### Auxiliary learning layer

Each data modality from the input layer goes through the biological dropconnect layer producing a set of output nodes of equal dimension (*C_GEX_*, *C_SNP_*). This layer aims to learn the latent space of one modality from the other. We consider a linear relationship between the two latent spaces, computed using **Equation 5**.

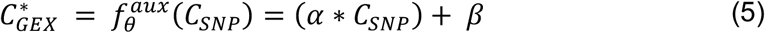

where *α and β* are scalar units representing weight and bias. We then concatenate the two latent space vectors and send them to a feed-forward neural network. One can get an average signal of the latent space vectors, but we decide not to pursue it as each latent node can be activated from both inputs or only one input.

#### Feed-forward classification layer

The concatenated gene layer is given to a fully connected feed-forward neural network with multiple hidden layers where each neuron in the hidden layers receives inputs from all previous layer outputs. ReLU activation is applied that forces negative weights to zero and handles non-linearity. ReLU activation is defined as shown in **Equation 6**.

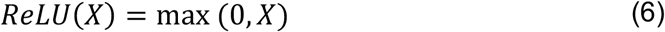

The final hidden layer is given to a softmax function to predict the output classes from the input given by **Equation 7**.

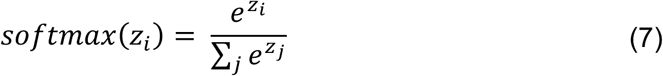

where *z* represents the neuron values from the previous layer.

#### Training of DeepGAMI model

Training the DeepGAMI model involved minimizing the loss function, combining primary task (phenotype prediction) loss and secondary task (cross-modality estimation) loss. The loss function used for the primary task is the cross-entropy loss (**Equation 8**) and mean squared error (MSE) loss is used for the secondary task (**Equation 9**).

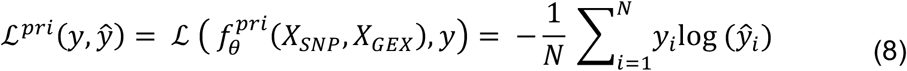

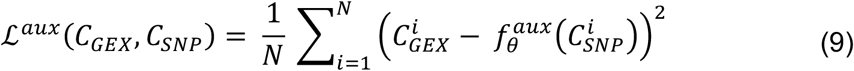

The overall objective loss function for the model is computed using **Equation 10**.

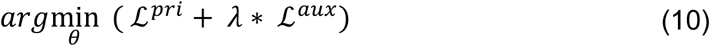

where *i* represents the *i^th^* training sample, *θ* is a set of parameters. *y and ŷ* represents the ground truth and the predicted labels respectively. We train the model with Adam Optimizer^104^, a gradient descent variant. The model is run over 100 epochs employing early stopping to avoid overfitting.

The data was split into training (80%) and test (20%) sets. The training was done using five-fold cross-validation (CV). Feature selection and five-fold cross-validation was applied on the training set to avoid information leak. All hyperparameters were selected based on the performance on five-fold CV performance. A grid search was used for choosing the optimal hyperparameters. A complete list of parameters used is available in **Supplementary Table 1**. After selecting the model with optimal hyperparameter combination, it was then used to test the performance on the held-out test set. As traditional accuracy measure can be misleading on the skewed datasets, we decided to use balanced accuracy (BACC) and area under the receiver operating characteristic curve (AUC) to evaluate performance. DeepGAMI is coded in pytorch^105^.

### Feature prioritization

After DeepGAMI is fully trained, thereby learning the optimal parameters, we use Integrated Gradient (IG)^106^ for prioritizing the input nodes (SNPs, TFs) as well as the latent space nodes (genes) along with their connection importance. The underlying mechanism of IG is that it computes the gradient for the model prediction for each input and measures the change in the output response based on the small changes in the input. This helps us detect SNPs and TFs attributed to the phenotype’s outcomes and can provide potential clues in understanding the underlying relationships (SNPs to TGs and TFs to TGs) for the given phenotype. IG is implemented using the Captum^107^ package in Python. We also used IG to prioritize the links between the input and the intermediate layer (SNPs and TFs to genes). Using the link importance score, we can fine-tune the regulatory networks for different phenotypes.

### Enrichment analysis

From the prioritized functional links between SNPs and TFs to genes, we extract the genes having the most important links (top 10% of the link importance scores). We then perform enrichment analysis on these genes using Metscape^88^. We used all genes used in our analysis as backgrounds for each data source. Meaningful biological enrichments (binomial FDR p-value < 0.05) were assigned as the predicted functionality of the group of prioritized genes.

### Multimodal data sources and processing

We applied DeepGAMI on several multimodal datasets with different phenotypes in human and mouse brains.

#### Schizophrenia cohort dataset

We use the population level bulk RNA-seq and genotype data for the human dorsolateral prefrontal cortex (DLPFC) from the psychENCODE consortium^46^ for predicting SCZ versus healthy individuals. RNA-seq data consists of normalized gene expression of 14906 genes for 1818 individuals. The genotype data include dosage information of more than 23 million SNPs for 672 individuals. eQTLs were downloaded from the brain eQTL meta-analysis study^22^ and we applied the scGRNom^108^ pipeline to extract GRN. We only included SNPs and genes which were present in eQTLs and GRNs. We then used t-test for feature selection as we had the curse of dimensionality issue. We ended up with 2080 SNPs, 126 TFs, and 84 intermediate layer genes for 343 control and 275 SCZ individuals.

We also tested DeepGAMI on the genotype and cell-type specific gene expression data from the CommonMind Consortium imputed using bMIND^92^ and a reference panel of 4 cell populations: GABAergic (i.e. inhibitory) neurons, glutamatergic (i.e. excitatory) neurons, oligodendrocytes, and a remaining group composed mainly of microglia and astrocytes. The reference panel for each cell population was constructed by taking the mean log2 counts per million for each gene across 32 brain donors^109^. With the prior information from this reference panel, bMIND adopts a Bayesian approach to impute the cell-type specific expression of each gene in each bulk sample from gene expression assayed from brain homogenate. DeepGAMI was applied following the same processing steps as mentioned above. We used cell-type specific eQTLs^22^ and applied scGRNom to each cell type for GRNs. **Supplementary Table 2** summarizes the number of features used as input to the model for these four cell types.

#### Alzheimer’s disease cohort dataset

ROSMAP cohort contains multi-omics data of the DLPFC brain region for aging and Alzheimer’s disease among humans^50^. We use the bulk RNA-seq and genotype data from this cohort for our analysis. RNA-seq data contains the FPKM gene expression values for 582 samples, and the genotype data contains the dosage information for 1709 individuals. For the DropConnect layer, we require GRNs and eQTLs to be given as priors. We extracted the eQTL information from the GTEx consortium^70^ for the human brain frontal cortex (BA9), which contains 146,763 eQTLs, and used the GRN from the psychENCODE consortium^46^. Clinical phenotypes included in our analysis are cognitive diagnosis (COGDX) score (that gives the cognitive status) ranging between 0 to 6, CERAD score (semi-quantitative measurement of the neuritic plaques useful for determining AD) ranged 0-4, and BRAAK score (semi-quantitative measurement for neurofibrillary tangle pathology) containing six stages. As the number of samples for scores within each range in each of these clinical phenotypes was very low, we coded the BRAAK phenotype into two classes (early-stage AD which contains braak stages of 0-3 and late-stage AD containing braak stages of 4-6), CERAD scores into three classes (No AD with scores 3-4, AD probable with score 2, and AD definite with score 0-1), and COGDX into three categories (No cognitive impairment (CI) with scores 0-1, mild CI with scores 2-3, and CI(AD/Dementia) with scores 4-6) using the coding available in ROSMAP. We used ANOVA to filter out SNPs and genes with high variance except for BRAAK, where we used a t-test instead of ANOVA for each clinical phenotype. We then filtered the SNPs and genes to match with eQTLs and GRN. The final number of features and class labels are shown in **Supplementary Table 3**. We ended up with 229 early-stage AD individuals and 275 late-stage AD individuals for the BRAAK score phenotype. For the COGDX phenotype, we had No CI (n=166), Mild CI (n=130), and CI(AD/Dementia, n=208) individuals. Finally, for the CERAD phenotype, we ended up with No AD (n=184), AD probable (n=171), and AD definite (n=149) individuals.

#### Mouse visual cortex

Patch-seq dataset contains single-cell multimodal data for the visual cortex brain region in mouse species for neuronal cells. This dataset includes transcriptomics and electrophysiological (ephys) data for 4,435 cells, where the electrophysiological data contain responses to three stimuli totaling up to 47 features^58^. We used the cell layers that reveal the location of the cells in the visual cortex as the cellular phenotype. The cell layer phenotype involves five layers (L1, L2/3, L4, L5, and L6). We followed the data extraction and preprocessing as done in DeepManReg^83^ and ended up with 41 ephys features and 1000 genes for 3,654 cells. We also used 112 layer4 (L4) neuronal cells Patch-seq data containing gene expressions but only a small set of electrophysiological data for independent testing^90^.

For this dataset, the inputs to the model are gene expression matrix comprised of m genes and k samples and is represented by *X_GEX_* ∈ *R^k*m^*, and electrophysiological features denoted by *X_ephys_* ∈ *R^k*n^*(*k* samples by *n* ephys features). The model is trained to optimize the parameters based on the modified loss functions from **Equations 8 and 9**, and the updated loss function is shown in **Equations 11 and 12**.

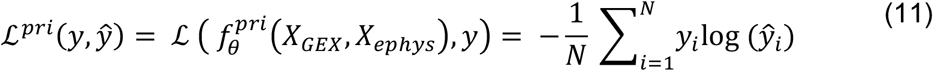

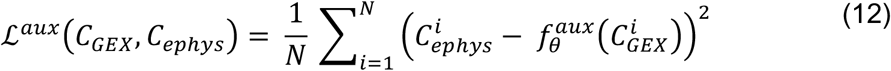

The overall objective loss function for the model is computed using **Equation 13**.

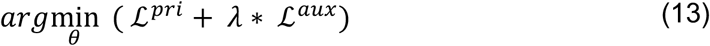

## Acknowledgments

The authors wish to thank all members of the Wang lab for insightful discussions on the topic.

## Data availability

The PsychENCODE bulk gene expression file for schizophrenia disorder can be downloaded from http://resource.psychencode.org/Datasets/Derived/DER-01_PEC_Gene_expression_matrix_normalized.txt, and the genotype data can be accessed from http://resource.psychencode.org. The cell type specific reference panel used for gene expression imputation is available at https://www.synapse.org/#!Synapse:syn22321061, and the imputed gene expression is available at https://www.synapse.org/#!Synapse:syn23234712. Alzheimer’s disease cohort data is available at https://www.synapse.org/#!Synapse:syn23446022. The processed gene expression and electrophysiological data from Patch-seq in mouse visual cortex is available at https://github.com/daifengwanglab/scMNC.

## Code availability

All code was implemented in Python using PyTorch as the deep learning package and the source code is publicly available at https://github.com/daifengwanglab/DeepGAMI.

## Supplementary information

- Data S1: Prioritized bulk genes and SNPs for schizophrenia.
- Data S2: Prioritized bulk transcription factors to genes and SNPs to gene links for schizophrenia.
- Data S3: Prioritized transcription factors to genes and SNPs to gene links for cognitive impairment phenotype in Alzheimer’s disease
- Data S4: Prioritized genes and electrophysiological features for cell cortical layers in mouse visual cortex.
- Data S5: Prioritized genes and SNPs for cognitive impairment phenotype in Alzheimer’s disease.
- Data S6: Prioritized cell-type genes and SNPs for schizophrenia.
- Data S7: Prioritized cell-type transcription factors to genes and SNPs to gene links for schizophrenia.
- Supplementary Materials: Supplementary Tables: 1-6 and Supplementary Figure 1.

## Abbreviations

GRN: Gene regulatory network
eQTL: Expression quantitative trait loci
GWAS: Genome-wide association studies
SCZ: Schizophrenia
AD: Alzheimer’s disease
CI: Cognitive impairment
MLP: Multilayer perceptron
SNP: Single nucleotide polymorphism

## Author Contributions

D.W. conceived the study. D.W. and P.B.C. designed the methodology and experiments. P.B.C., C.H., T.J. J.B., and J.F. curated and processed data required for the analysis. J.W. and G.H. imputed cell-type gene expression data of individuals. P.B.C., C.H., T.J., and S.K. performed analysis and visualization. P.B.C. and S.A. implemented the software. P.B.C., J.W., G.H., P.R. and D.W. edited and wrote the manuscript. All authors read and approved the final manuscript.

