## Supplementary Tables: 1-6 and Supplementary Figure 1 for "DeepGAMI: Deep biologically guided auxiliary learning for multimodal integration and imputation to improve phenotype prediction"

Deep auxiliary learning for multi-modal integration and estimation to improve genotype-phenotype prediction

| **Hyper-parameters** | **Values** |
| --- | --- |
| Number of latent dimensions | {250, 500, 1000} |
| Number of hidden layers | {1, 2, 3} |
| Number of neurons | {50, 100, 250, 500, 1000} |
| Dropout rate | {0.25, 0.5, 0.75} |
| L2 regularization rate | {0.0001, 0.001, 0.01, 0.1} |
| Learning rate | {0.0001, 0.001, 0.01, 0.1} |
| λ (Auxiliary loss regularization) | {0.25, 0.5, 1} |

**Supplementary Table 1 - List of all hyperparameters used in DeepGAMI**

**Supplementary Table 2 - Summary table showing features and class labels for different available phenotypes for ROSMAP AD dataset.**

| **Phenotype** | **TFs** | **SNPs** | **Genes** | **Class Labels** |
| --- | --- | --- | --- | --- |
| COGDX score | 102 | 273 | 354 | No CI, Mild CI, and CI (AD/dementia) |
| CERAD score | 98 | 467 | 369 | No AD, AD probable, and AD definite |
| BRAAK staging | 114 | 544 | 366 | Early stage and late stage |

**Supplementary Table 3 – Balanced accuracy comparison for ROSMAP AD dataset.**  The table compares performance of DeepGAMI with other baseline ML methods for three different phenotypes: CERAD score, COGDX score, and BRAAK staging

|  | **CERAD score (3 classes)** | **COGDX score (3 classes)** | **BRAAK Staging (binary)** |
| --- | --- | --- | --- |
| Random Forest | 0.35 | 0.383 | 0.538 |
| Naïve Bayes | 0.445 | 0.34 | 0.555 |
| MLP | 0.378 | 0.385 | 0.742 |
| DeepGami Dual | 0.681 | 0.688 | 0.806 |
| DeepGami Single | 0.68 | 0.682 | 0.79 |

**Supplementary Table 4 – Balanced accuracy comparison for Patch-seq dataset.**  The table compares five-fold cross-validation balanced accuracies of DeepGAMI with other several ML methods across five cell layers for visual cortex region in mouse brain.

|  | **L1** | **L2/3** | **L4** | **L5** | **L6** |
| --- | --- | --- | --- | --- | --- |
| **Unregularized** | 0.308 ± 0.177 | 0.312 ± 0.155 | 0.314 ± 0.169 | 0.302 ± 0.157 | 0.3 ± 0.155 |
| **DeepManReg** | 0.515 ± 0.071 | 0.518 ± 0.031 | 0.514 ± 0.055 | 0.511 ± 0.034 | 0.511 ± 0.047 |
| **LMA** | 0.439 ± 0.093 | 0.431 ± 0.051 | 0.435 ± 0.073 | 0.425 ± 0.05 | 0.428 ± 0.059 |
| **CCA** | 0.467 ± 0.079 | 0.462 ± 0.042 | 0.466 ± 0.053 | 0.459 ± 0.041 | 0.462 ± 0.056 |
| **Matcher** | 0.471 ± 0.086 | 0.466 ± 0.032 | 0.466 ± 0.055 | 0.465 ± 0.041 | 0.463 ± 0.046 |
| **DeepGamiDual** | 0.75 ± 0.076 | 0.689 ± 0.067 | 0.564 ± 0.095 | 0.484 ± 0.085 | 0.797 ± 0.075 |
| **DeepGamiSingle** | 0.765 ± 0.073 | 0.648 ± 0.069 | 0.61 ± 0.094 | 0.443 ± 0.083 | 0.815 ± 0.068 |

**Supplementary Table 5 – Summary table of the features for cell-type-specific Schizophrenia dataset**

| **Cell type** | **TFs** | **SNPs** | **Genes** |
| --- | --- | --- | --- |
| Oligodendrocytes | 247 | 339 | 66 |
| Microglia | 552 | 231 | 108 |
| Inhibitory neurons | 465 | 206 | 88 |
| Excitatory neurons | 414 | 198 | 73 |

**Supplementary Table 6 – Binary Classification results for cell-type-specific Schizophrenia dataset.**  The table shows five-fold cross validation balanced accuracy (BACC) comparison of DeepGAMI against several machine learning algorithms. Each cell represents average BACC along with standard deviation.

|  | **Microglia** | **Oligodendrocytes** | **Inhibitory Neurons** | **Excitatory Neurons** |
| --- | --- | --- | --- | --- |
| Random Forest | 0.563 ± 0.049 | 0.586 ± 0.036 | 0.576 ± 0.026 | 0.5474 ± 0.047 |
| Naïve Bayes | 0.633 ± 0.05 | 0.634 ± 0.042 | 0.621 ± 0.083 | 0.6469 ± 0.048 |
| MLP | 0.743 ± 0.058 | 0.661 ± 0.023 | 0.715 ± 0.051 | 0.6828 ± 0.051 |
| Varmole | 0.765 ± 0.026 | 0.733 ± 0.053 | 0.741 ± 0.02 | 0.724 ± 0.019 |
| DeepGami Dual | 0.795 ± 0.035 | 0.762 ± 0.053 | 0.755 ± 0.027 | 0.758 ± 0.022 |
| DeepGami Single | 0.784 ± 0.024 | 0.759 ± 0.056 | 0.745 ± 0.036 | 0.746 ± 0.034 |

**Supplementary Figure 1 – Integrated Gradient results for Patch-seq mouse visual cortex data.** Corrplot comparing the importance score of all 41 electrophysiological features across the five cell layers derived from DeepGAMI.

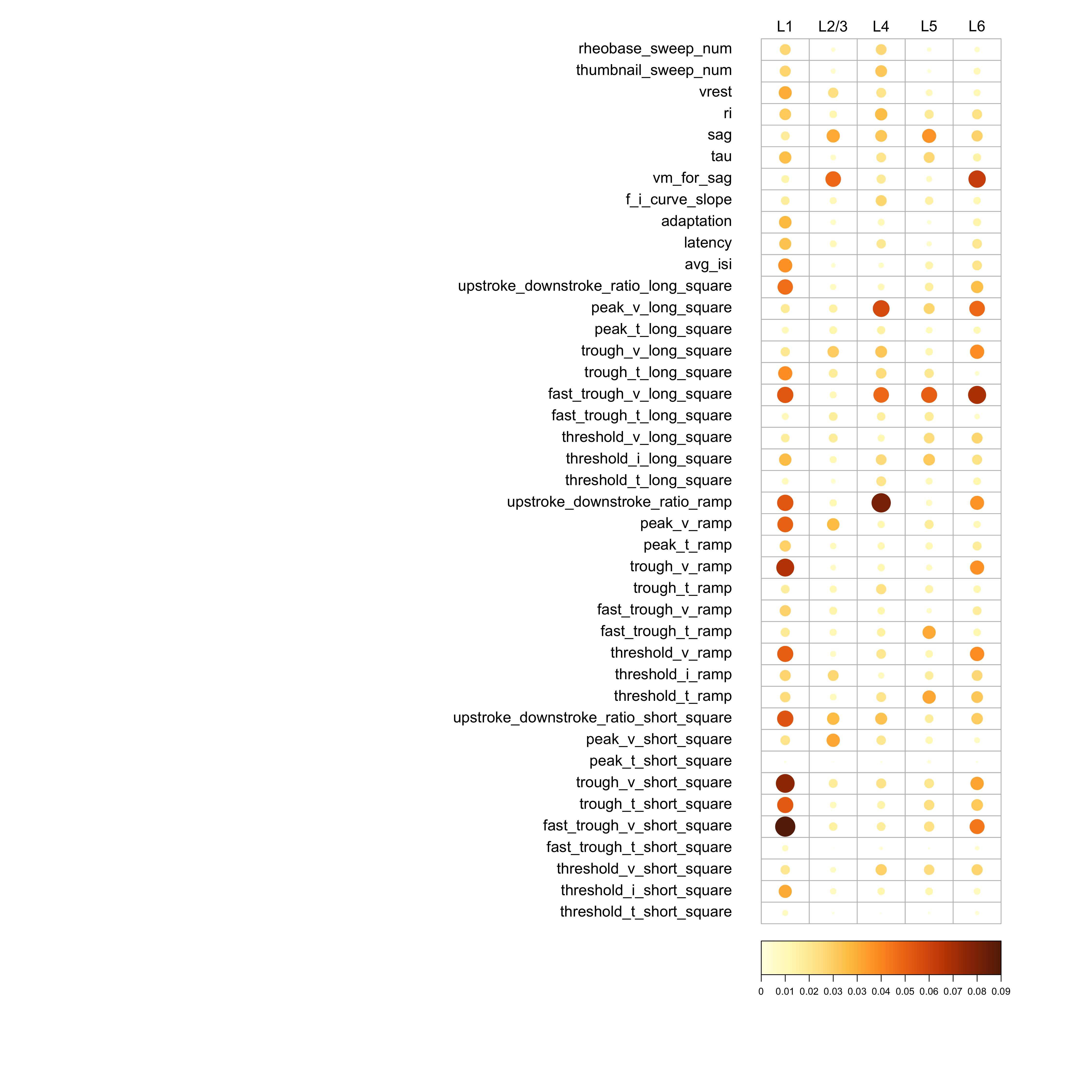

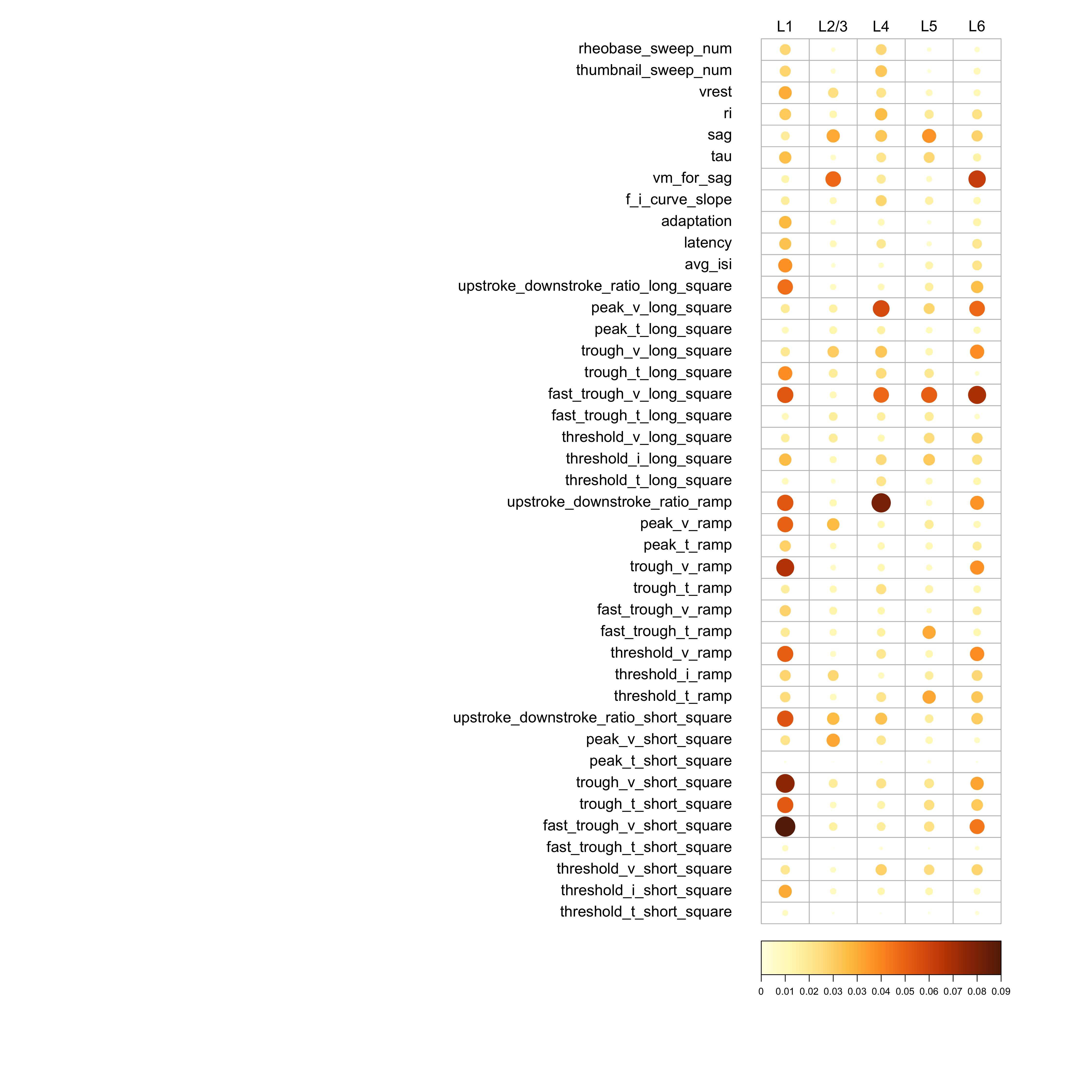

Importance score
